# Differential Cetacea Circadian Rhythmicity is associated with the molecular erosion of Cortistatin

**DOI:** 10.1101/2020.05.11.087833

**Authors:** Raul Valente, Luís Q. Alves, Matilde Nabais, Filipe Alves, Isabel Sousa-Pinto, Raquel Ruivo, L. Filipe C. Castro

**Author notes:** Corresponding author: L. Filipe C. Castro and Raquel Ruivo, CIIMAR/CIMAR U. Porto, Av. General Norton de Matos s/n, 4450-208 Matosinhos, Portugal. These authors contributed equally (Raul Valente, Luís Q. Alves).

## Abstract

The ancestors of Cetacea underwent profound morpho-physiological alterations. By displaying an exclusive aquatic existence, cetaceans evolved unique patterns of locomotor activity, vigilant behaviour, thermoregulation and circadian rhythmicity. Deciphering the molecular landscape governing many of these adaptations is key to understand the evolution of phenotypes. Here, we investigate Cortistatin (*CORT*), a neuropeptide displaying an important role mammalian biorhythm regulation. This neuropeptide is a known neuroendocrine factor, stimulating slow-wave sleep, but also involved in the regulation of energy metabolism and hypomotility inducement. We assessed the functional status of *CORT* in 139 mammalian genomes (25 orders), including 30 species of Cetacea. Our findings indicate that cetaceans and other mammals with atypical biorhythms, thermal constraints and/or energy metabolism, have accumulated deleterious mutations in *CORT*. In light of the pleiotropic action of this neuropeptide, we suggest that this inactivation contributed to a plethora of phenotypic adjustments to accommodate adaptive solutions to specific ecological niches.

## Main text

Habitat transitions nourish the emergence of novel phenotypes. In this context, Cetacea (whales and dolphins) are a particularly fascinating group to understand the genomic signatures of adaptation to radical shifts, given their land-to-water evolutionary history and their exclusive reliance on aquatic ecosystems (e.g. McGowen et al. 2014; Huelsmann et al. 2019; McGowen et al. 2020). Specifically, gene loss events, including complete gene absence or sequence gene erosion (Albalat and Cañestro, 2016), seem to stand out as significant triggers of phenotypic adaptations within this group: as reported for disparate processes such as skin remodelling, immunity and inflammation, deep-diving induced hypoxia, blood pressure maintenance, or even circadian rhythmicity and sleep/vigilance behaviours (e.g. Braun et al. 2015; Hecker et al. 2017; Lopes-Marques et al. 2018, 2019a, 2019b, 2019c; Alves et al. 2019; Ehrlich et al., 2019; Huelsmann et al. 2019). Among these, circadian and sleep/vigilance behaviours are particularly challenging since cetaceans require occasional surfacing to breathe. Importantly, mammalian sleep generally prompts several physiological adjustments, which conflict with a fully aquatic lifestyle, including hypomotility, sleep thermoregulation or decreased blood pressure (Giglio et al. 2007; Alves et al. 2019). To offset these constraints, Cetacea exhibit distinctive biological rhythms, allowing the maintenance of vigilant states over long periods of time (Ridgway et al. 2006; Branstetter et al. 2012), along with lateralized sleep (Lyamin et al. 2008), or even uninterrupted activity as observed in Delphinidae neonates and mothers (Lyamin et al. 2005; Siegel 2005). In agreement, recent studies have highlighted gene loss signatures related with the maintenance of such an unusual form of mammalian sleep/vigilance behaviours, notably regarding the synthesis and signalling of melatonin, a potent modulator of circadian rhythmicity, affecting multiple physiological and behavioural processes such as sleep entrainment, locomotor activity or thermoregulation (Huelsmann et al. 2019; Lopes-Marques 2019c). Here, we investigate Cortistatin (*CORT*), a cyclic neuropeptide which plays an important role in sleep physiology (de Lecea et al. 1996). Belonging to the somatostatin (SST) neuropeptide family, *CORT* was shown to have sleep-promoting properties, stimulating slow-wave sleep, as well as to induce hypomotility (Spier and de Lecea 2000) (Figure 1). Moreover, the diversification of roles attributed to *CORT* also includes regulation of endocrine metabolism, immunomodulation, inflammatory responses, pain perception and cardiovascular protection (Deghengi et al. 2001; Robas et al. 2003; Broglio et al. 2007; Gonzalez-Rey et al. 2015; Liang et al. 2019). Despite sharing similarities with SST, such as peptide structure, resulting from proteolytic cleavage, and binding affinity towards SST receptors, *CORT* generally yields antagonizing effects when compared to SST (Spier and de Lecea 2000; Broglio et al. 2007). In this context, we sought to characterize the functional status of *CORT* genes in Cetacea, and other mammalian lineages, to determine whether gene inactivation events have taken place, as previously reported for other genes (e.g. Kim et al. 2014; Shinde et al. 2019). We began by inspecting the open reading frame (ORF) of *CORT* in a selected sub-set of Cetacea species using PseudoIndex (Alves et al. 2020). This user assistant metric built into the Pseudo*Checker* pipeline rapidly estimates the erosion condition of the tested genes - discrete scale from 0 (functional) to 5 (pseudogenized) (Alves et al. 2020). All analysed species revealed a PseudoIndex equal to 5 within Odontoceti species, except for *Lipotes vexillifer* and *Delphinapterus leucas*, which displayed a PseudoIndex of 2 and 3 respectively (Supplementary File 1). This analysis suggested that the ORF of *CORT* includes inactivating mutations. Thus, we next performed a manual and exhaustive *CORT* sequence analysis in a larger phylogenetic collection of species and included an additional curated validation step of the mutational evidence. In detail, genomic *CORT* sequences from 139 terrestrial and aquatic mammalian species (Supplementary File 2), containing 30 Cetacea (9 Mysticeti and 21 Odontoceti), were collected from available genomes and manually annotated using *Bos taurus* (phylogenetically related to Cetacea (McGowen et al. 2019)) or *Homo sapiens* (for other mammals) *CORT* coding sequences (CDS) as a reference (Lopes-Marques et al. 2017). Predicted ORFs were screened for disrupting mutations, with these being further validated (when possible) using data from at least 2 independent Sequence Read Archive (SRA) genomic projects or from 2 distinct individuals (Lopes-Marques et al. 2019a). Gene orthology between Cetacea species and *B. taurus* was validated through synteny analysis, which essentially revealed *locus* conservation between species (Supplementary File 3).

**Figure 1:**
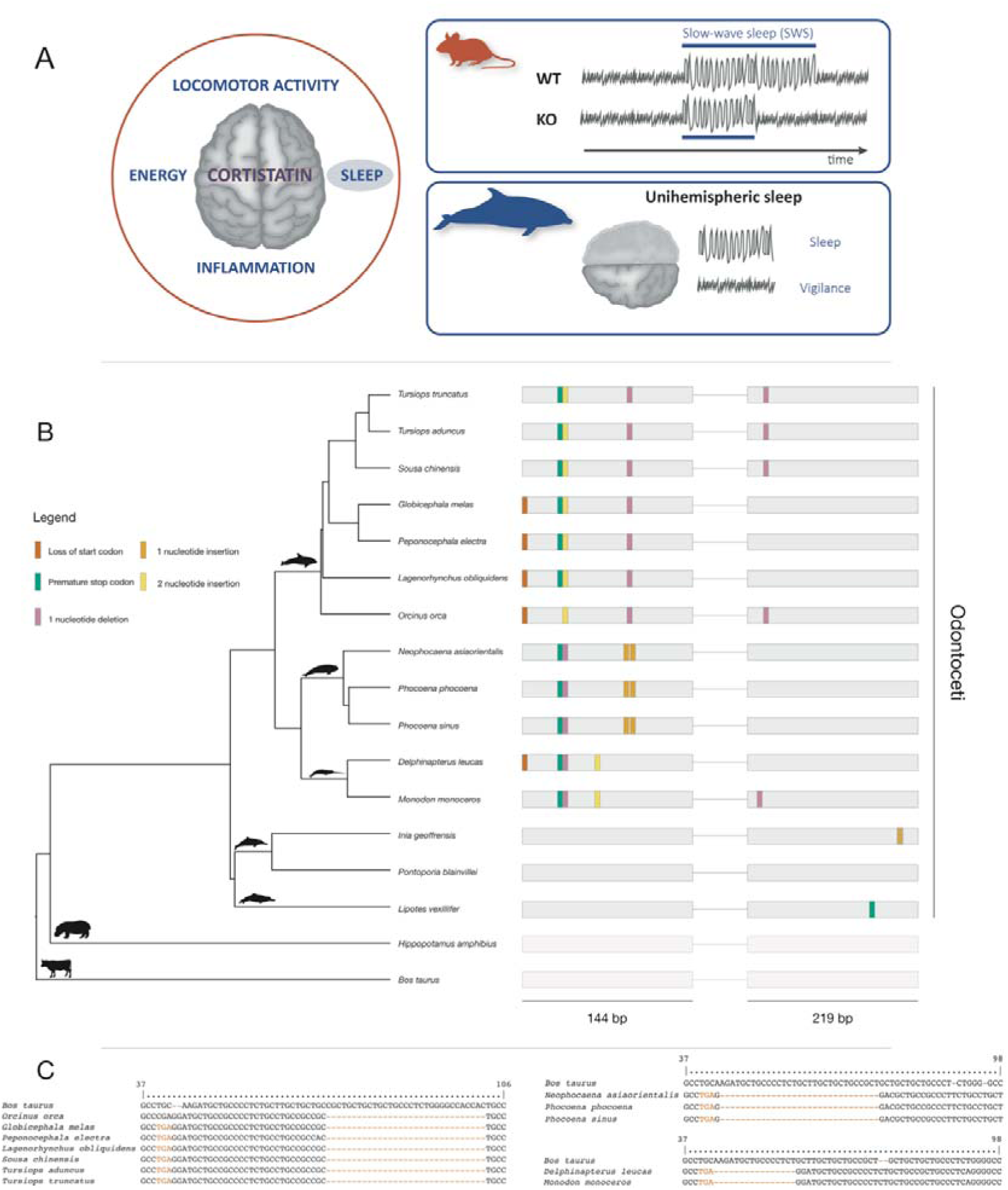
Distinct physiological roles attributed to *CORT*, effect of *CORT* knockout mice in slow wave sleep time and cetacean idiosyncratic sleeping in the form of lateralized sleep behaviour (A). Mutational landscape of the *CORT* gene in several Odontoceti, *H. amphibius* and *B. taurus* and location of identified mutations (B). Phylogenetic relationships are derived from McGowen et al. (2020). Example of *CORT* open reading frame (ORF) inactivating mutations concerning *Delphinidae, Phocoenidae* and *Monodontidae* clades (C). Numbers above characters represent the alignment position index. Silhouettes were recovered from Phylopic (http://phylopic.org).

### *CORT* exhibits a conserved premature stop codon in Delphinoidea

In Odontoceti (toothed whales and dolphins), all Delphinoidea, including *Delphinidae* (dolphins, e.g. *Tursiops truncatus, Orcinus orca*), *Phocoenidae* (porpoises, e.g. *Phocoena phocoena*), and *Monodontidae* (narwhal, *Monodon monoceros*, and beluga whale, *Delphinapterus leucas*), exhibited a conserved stop codon mutation in exon 1 (Figure 1), with the exception of *O. orca*, which displays an arginine codon in the same position (CGA). Yet, given the striking conservation of the premature stop codon across analysed Delphinoidea, and the single nucleotide difference between both codons, the *Orca* exception likely represents a case of mutational reversion (Stop>Arg, <u>T</u>GA><u>C</u>GA) (Rosenberg 2001). Nonetheless, in addition to the premature stop codon, other mutations were identified, confirming the erosion of *CORT* in *O. orca*, such as frameshifts in exon 1 (31 nucleotide deletion and 2 nucleotide insertion), conserved in all *Delphinidae* (Figure 1). In *O. orca* and Delphininae species (*Tursiops* and *Sousa*), exon 2 also presented a frameshift mutation (1 nucleotide deletion), a pattern not consistent with the species branching tree topology (McGowen et al., 2019). Additionally, deleterious mutations were also found across all members of *Phocoenidae* and *Monodontidae* (described in Supplementary File 4). Detected mutations were further validated by independent SRAs in all species, when available (Supplementary File 5).

To further scrutinize the functional condition of *CORT*, we searched for transcriptional evidence of this gene in *Globicephala melas, O. orca* and *D. leucas* by investigating available brain RNA-Seq projects at the SRA database (Krüger et al. 2020). The collected mRNA reads were mapped against the corresponding annotated gene and classified as spliced reads (reads partially covering two exons), exon-intron reads (unspliced reads) and exonic reads (reads containing only data from a single exon) (Lopes-Marques et al. 2019b). Briefly, we observed a substantially high proportion of exon-intron reads *versus* spliced reads in stark contrast to the pattern found in cow (positive control) (Figure 2). In addition, we were able to identify at least one premature stop codon in the transcripts of all analysed species (Supplementary file 6), further validating the predicted ORF-abolishing mutations.

**Figure 2:**
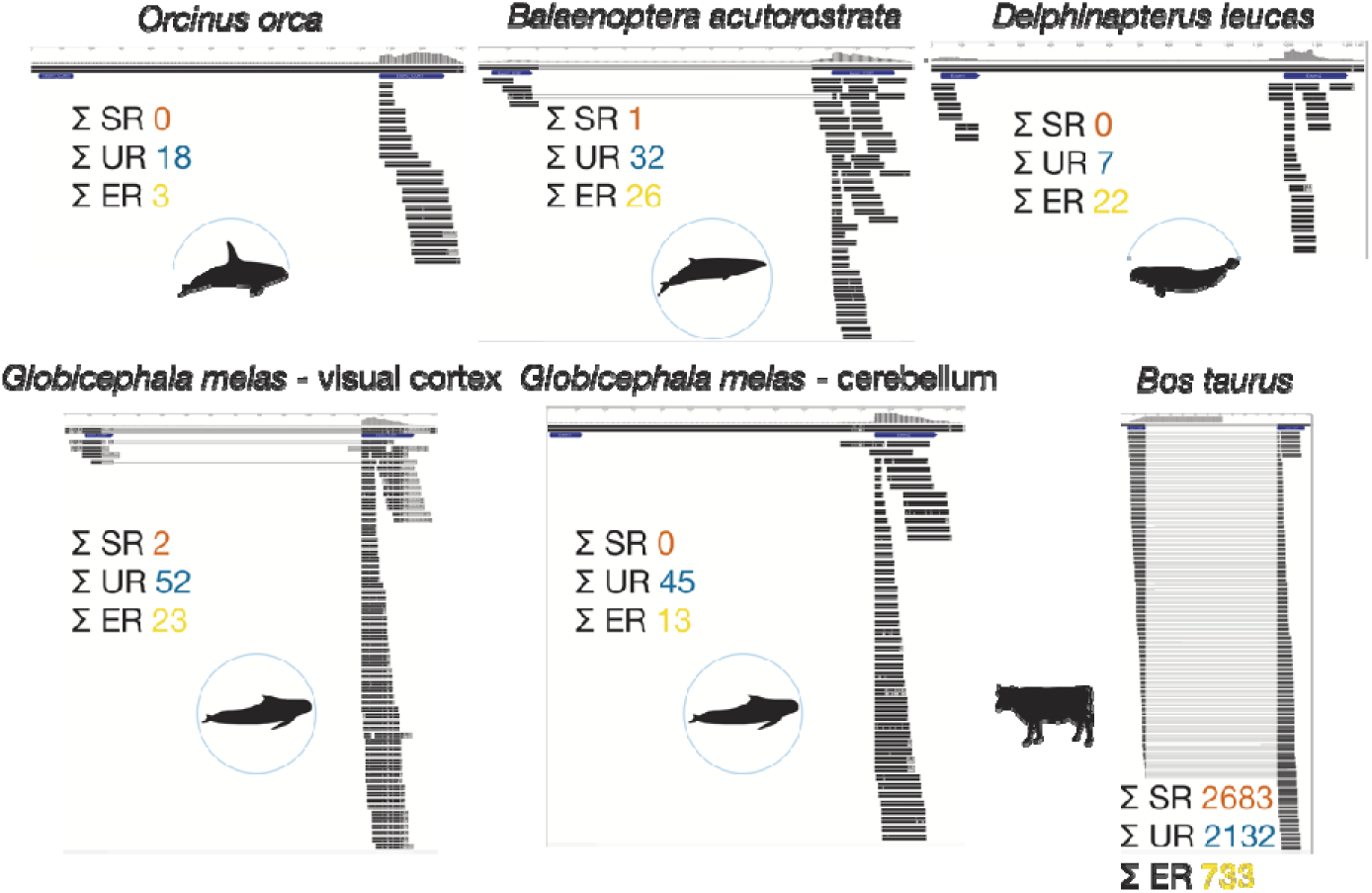
Gene expression of *CORT* across represented Cetacea species: mapping of the NCBI Sequence Read Archive (SRA) brain RNA-Seq reads (black) for each of the 5 represented species against the corresponding *CORT* annotated gene (in blue). Overall count of RNA-Seq mapped reads for each species is also represented. Reads are classified into spliced reads (SR), exon-intron reads (UR), and exonic reads (ER).

### Divergent patterns of functional inactivation are observed in river dolphins

River dolphins represent a polyphyletic group within Odontoceti, including species such as *Lipotes vexillifer, Pontoporia blainvillei, Inia geoffrensis* or *Platanista gangetica*. Among these, *L. vexillifer* showed a premature stop codon in exon 2, *P. gangetica* presented frameshift mutations in both exons (31 nucleotide deletion in exon 1 and 4 nucleotide insertion in exon 2) and *I. geoffrensis* displays an indel in exon 2 (Suplementary File 4). Given that no SRAs are available for these species, SRA validation was not performed. Regarding *P. blainvillei*, on the other hand, no ORF disrupting mutations were found. Yet, further sequence analysis revealed amino acid changes that possibly impair canonical peptide function (Supplementary File 7). *CORT*, similarly to SST, contains a carboxyl-terminal FWKT (Phe-Trp-Lys-Thr) tetramer, essential for binding to SST receptors (de Lecea et al. 1996; Spier and de Lecea 2000). Despite its strict conservation across mammals, amino acid substitutions are observed in *P. blainvillei*, thus suggesting that, in this species, SST receptor binding is possibly impaired (Supplementary File 7).

### Variable disruption patterns are observed in the Odontoceti Ziphiidae, Physeteroidea, as well as in Mysticeti

In *Ziphiidae* (*Mesoplodon bidens* and *Ziphius cavirostris*) or *Kogia breviceps* (pygmy sperm whale), no ORF disrupting mutations, nor radical amino acid replacements, were detected (Supplementary File 4). On the other hand, *Physeter macrocephalus* presented a single frameshift mutation in exon 2, validated by SRA (Supplementary file 4). In Mysticeti, *Balaenoptera acutorostrata* presented a premature stop codon in exon 2 (Supplementary File 4). Also, a 22-nucleotide deletion was detected in exon 1 of *Eschrichtius robustus, Megaptera novaeangliae* and *Balaenoptera musculus*, while for *Balaenoptera physalus* we identified three indel mutations also in exon 1. These mutations were validated by SRA searches in *B. acutorostrata* and *E. robustus* (Supplementary File 5). Besides these ORF-disrupting mutations, a conserved loss of start codon (ACG - Thr) was also detected for the full set of examined Mysticeti species, except for *B. acutorostrata*. However, an ATG (methionine) codon could be found downstream from the threonine in the same translational reading frame (Supplementary File 4). Thus, in species presenting no more disruptive mutations - namely in *Eubalaena glacialis, Eubalaena japonica, Balaenoptera bonaerensis* and *Balaenoptera edeni*, we cannot rule out the possibility of a functional *CORT* gene. In *Balaena mysticetus*, the fragmentation of the genomic region (Ns) corresponding to exon 2 from *CORT* impeded us to infer the coding status in this species. Additionally, for *B. acutorostrata*, mapping of transcriptional reads from a brain RNA-Seq project yielded a high proportion of exon-intron reads versus spliced reads (Figure 2) and further corroborated the presence of the premature stop codon in exon 2. For the remaining analysed species, no deleterious mutations were unequivocally found. Nonetheless, besides ORF disruptive mutations, other processes can lead to the non-functionalization of a gene: i.e. lack of transcription, RNA decay, or even protein degradation (e.g. Sadier et al. 2018). Yet, we were unable to verify the transcriptomic profile of *CORT* in these species given the absence of adequate transcriptomic data.

### Selection analysis

To find evidence of some relaxation of purifying selection typically associated with events of gene pseudogenization, RELAX analysis using the HyPhy package was performed (Pond et al. 2005; Wertheim et al. 2014). We used the predicted *CORT* sequences from 43 species, comprising all Cetartiodactyla species analysed in this study (Supplementary File 2). As RELAX compares a background set of species with a foreground set of species over a hypothesis-testing framework, we targeted the ancestral branch of the cetacean lineages, using the branches from all non-cetacean species as a reference. Although no significant intensification or relaxation has been found (Supplementary File 8 – Table S1), it can be seen through the general descriptive model (Supplementary File 8 – Figure S1) that the ancestor of all cetaceans experienced relaxed selection (k□<□1), coinciding with substantial changes in *CORT* amino acidic composition. Moreover, we detected signs of intermittent relaxed selection (k□<□1) within various cetacean lineages, which could result from differences in gene length/composition across branches. This could be due to an intensification of positive/diversifying selection prior to the observed pseudogenization in the terminal cetacean branches.

### Episodes of *CORT* ORF-disruption are found in other mammal species

We next interrogated the genomes of other mammalian species. The implemented assessment revealed a number of ORF-disrupting mutations in 14 species (Supplementary File 9). For example, the sequence of *CORT* in *Ursus maritimus* (polar bear) revealed a single nucleotide deletion in exon 2 and in *Procavia capensis* (rock hyrax) a premature stop codon in exon 2 (confirmed by SRA search in *U. maritimus*; Supplementary File 10); in the Pholidota *Manis javanica* and *Manis pentadactyla* (pangolins), a shared frameshift mutation in exon 1 and a premature stop codon was identified (validated by SRA in *M. javanica*; Supplementary File 10). In the Carnivora *Canis lupus* (wolf) and *Lycaon pictus* (African wild dog), conserved single nucleotide insertions were found in exon 2, whereas *Felis catus* (cat) and *Acynonix jubatus* (cheetah), exhibit a conserved 2 nucleotide insertion in exon 1, all validated by independent sequencing reads. Other examples include *Dasypus novemcinctus* (nine banded armadillo) with an indel and a premature stop codon in exon 2, *Choleopus hoffmanni* (Hoffmann’s two-toed sloth) with single nucleotide insertions and deletions in both exons, and *Condylura cristata* (star-nosed mole) for which exon 2 was not found.

### Metabolic Homeostasis and Circadian rhythmicity

Our study provides clear evidence for the dismantling of *CORT* gene sequence within most Cetacea and in several other mammalian lineages. Curiously, various of these species display particular biological rhythms and/or labile body temperatures: the hibernating polar bear; the subterranean star-nosed mole; nocturnal species with variable and relatively low body temperatures such as the two-toed sloth, the nine banded armadillo, pangolins, or the rock hyrax that undergoes nocturnal hypothermia (Ralph 1975; Brown and Downs 2007; Ware et al. 2013). Thus, it is plausible to hypothesize that, in these species, *CORT* inactivation is related with changes in circadian rhythmicity, affecting sleep/vigilance behaviours, body temperature maintenance and/or activity patterns. Interestingly, these species also display some degree of reduction or even complete atrophy of the pineal gland, the dominant organ for melatonin synthesis (Pévet 2002). In Cetacea, constituting a group with a remarkably altered sleep physiology, *CORT* pseudogenization possibly paralleled additional losses, notably regarding the melatonin synthesis and signalling genes (Lopes-Marques et al. 2019c; Huelsmann et al. 2019). Unexpectedly, we also found robust evidence supporting *CORT* pseudogenization in some Carnivora species, particularly felids and canids. However, and in addition to SST receptors, *CORT* was also suggested to bind other G-protein-coupled receptors such as the ghrelin receptor growth hormone secretagogue receptor type 1a (GHSR-1a), which participates in energy homeostasis regulation: namely in the control of food intake and energy metabolism (Deghenghi et al. 2001; Broglio et al. 2007). Mice lacking *CORT* gene showed alterations in whole-body metabolism, in a gender-dependent manner, notably resulting in the increase of acylated ghrelin levels in female, or higher glucose levels and insulin resistance in males (Córdoba-Chacón et al. 2011; Luque et al. 2016). In addition, *CORT* action was also proposed to be conditioned by the underlying metabolic status (i.e. fasting or obesity) (Luque et al. 2016). These observations are in agreement with the metabolic profile observed in felids, associated with a low-carbohydrate consumption and deficit in hepatic glucokinase activity, yielding fasting hyperglycaemia and insulin resistance (Schermerhorn 2013). Both hyperglycaemia and insulin resistance were also reported in dolphins, also displaying reduced carbohydrate intake (Schermerhorn 2013). Overall, our results put forward that the evolutionary loss of *CORT*, along with additional genomic and phenotypic signatures, paralleled the adjustment of circadian rhythmicity and energy homeostasis to accommodate adaptations to specific ecological niches and life-history traits.

## Acknowledgments

We acknowledge the various genome consortiums for sequencing and assembling the genomes.

